# Septin Architecture Dictates Size-Dependent Diffusion Barriers on biomimetic Membranes

**DOI:** 10.1101/2025.10.08.681182

**Authors:** Brieuc Chauvin, Koyomi Nakazawa, Ingrid Adriaans, Marco Geymonat, Simonetta Piatti, Bassam Hajj, Stéphanie Mangenot, Aurélie Bertin

**Affiliations:** Laboratoire Physico Chimie Curie, Institut Curie, PSL Research University, Sorbonne Université, CNRS UMR168, 75005, Paris, France; Centre de Biologie Cellulaire de Montpellier (CRBM), Université de Montpellier - CNRS UMR 5237, Montpellier, France; Laboratoire NABI, Université Paris Cité, CNRS UMR 1875, 75006 Paris, France; Department of Genetics, University of Cambridge, Downing Street, Cambridge CB2 3EH, United Kingdom

## Abstract

Septins are cytoskeletal filaments bound to the inner leaflet of the plasma membrane, essential for cell division. They are involved in the formation of diffusion barriers for membrane-bound components. Whether septins directly function as barriers or as part of a broader regulatory cascade of events remains unclear. We addressed this using in vitro reconstituted assay of biomimetic synthetic membranes. We probed whether the diffusion of biomimetic fluorescent objects with tunable steric hindrance – mimicking membrane bound proteins – is constrained by septins. Our results indicate that (i) lipids and transmembrane proteins lacking cytosolic domains diffuse freely in the presence of septins (ii) membrane-bound objects with large cytosolic domains experience size-dependent diffusion constraints; and (iii) the ability of septins to act as diffusion barriers is finely tuned by their intrinsic filament organization. These findings demonstrate that septins can directly impose size-selective diffusion barriers and that their filament organization critically tunes this function.

## Introduction

Septins are cytoskeletal filamentous proteins that assemble into the inner leaflet of the plasma membrane and are implicated in the formation of diffusion barriers across various cell types^1^. In fungi, septins localize to the bud neck and are essential for maintaining asymmetry during cell division^2–4^. In mammals, they accumulate at compartment boundaries such as the base of cilia^5^, the annulus of spermatozoa^6^, and dendritic branch points^7^—sites where lateral diffusion of membrane components is restricted. These observations position septins as potential organizer of compartmentalization in membranes, but a key question remains: do septins create the barrier them self or they act as a part of a larger cascade of events?

Diffusion barriers, at the plasma membrane, endoplasmic reticulum and nuclear membranes are necessary to ensure cellular functions^1^, such as the establishment of polarity during cell division^8^, and directing cellular compartmentalization. For instance, diffusion barriers are essential in controlling lateral membrane diffusion at the base of cilia^5^, base of dendrites in neurons^7^, and at the annulus of spermatozoa^9^. Membrane diffusion is tightly regulated by the composition and organization of lipids and proteins, particularly through the formation of high-density membrane domains that interact with cytoskeletal components and the extracellular matrix. Shingolipids were also shown to be involved in diffusion barriers at the endoplasmic reticulum^10^.As a result, lipid diffusion in living cells is significantly slower than in reconstituted membranes, highlighting the importance of the cytosolic environment and associated scaffolding structures^11^.

Among the cytoskeletal players, actin was shown to affect and control the diffusion of lipids and membrane interacting proteins both in situ^12,13^ and in in vitro reconstituted experiments^14^. Actin filaments were demonstrated to restrict the diffusion of DOPE lipids in situ^11^ which diffuse in a so called “hop” diffusive mechanism. Septins have received less attention. In budding yeast, a model system for studying diffusion of transmembrane proteins, septins were shown to be required for the restricted mobility of transmembrane proteins such as Spa2 (from the polarisome), Sec3, and Sec5 (from the exocyst complex), which contain large cytosolic domains, as well as for protein sequestration between mother and daughter cells^8^. More recently, Pacheco et al.^15^ showed that septins, as opposed to other proteins, was able to corral PI(4,5)P2 using fluorescence microscopy assays in reconstituted supported lipid bilayers, suggesting a major role for septins in establishing direct physical diffusion barriers. Similarly, septin depletion disrupts compartmentalization of membrane-associated proteins in neurons^7^ and spermatozoa^9^. However, these in vivo studies cannot distinguish whether septins act as direct physical barriers or as regulators within larger molecular cascades.

Septins interact directly with phosphoinositide-rich membranes^16–18^, and in vitro studies have shown that both yeast and human septins can assemble into membrane-bound filaments. Interestingly, the supramolecular organization of these filaments differs by species: budding yeast septins form parallel arrays^18–20^, while human septins form orthogonal meshes^21^, suggesting possible differences in barrier-forming capability. In vitro, we and others could show that both budding yeast and human septins can reshape Giant Unilamellar Vesicles (GUVs) in a curvature-sensitive manner^19,21^. Septins also interact with other cytoskeletal elements, such as actin^22^ and microtubules^23,24^, as well as with regulatory partners including Bud3, Bud4, and Gic proteins^25–29^. These interactions govern septin organization and function in vivo, yet whether septins alone are sufficient to block lateral diffusion remains unknown.

To test whether septins can impose a physical barrier for membrane diffusion, we developed a minimal in vitro reconstituted assay using supported lipid bilayers (SLBs) and giant unilamellar vesicules (GUVs) in combination with fluorescently labeled molecules of varying sizes and membrane anchoring strategies. We used either Fluorescence Recovery After Photobleaching (FRAP) or single particle tracking (SPT) to quantify the diffusion in the presence and absence of septin assemblies. To avoid potential surface effects that may impact the results observed using supported lipid bilayers, we have mostly followed the diffusion of fluorescent object at the bottom of GUVs, focusing on free standing membranes. We compared the effects of budding yeast and human septins, which differ in their filament arrangements, to examine whether structural organization influences barrier strength. This reconstituted approach allowed us to isolate the direct physical contribution of septins to membrane diffusion barriers, free from cellular complexity.

## Results

### Septins minimally affect the diffusion of lipids

Septins are known to interact and self-organize on lipid membranes^18,19,21,30,31^, yet it remains unclear whether their membrane-bound filamentous structures would directly impact the diffusion of lipids. To address this we tested the impact of yeast septin filaments covalently tagged with GFP (Green Fluorescent Protein) on the diffusion of lipids in reconstituted membrane. We used supported lipid bilayers (SLB) composed of EggPC (56 %), Cholesterol (15 %), 1,2-di-(9Z-octadécénoyl)-sn-glycéro-3-phosphoéthanolamine (DOPE) (10 %), 1,2-dioleoyl-sn-glycero-3-phospho-L-serine (DOPS) (10 %) and L-α-phosphatidylinositol-4,5-bisphosphate (PI(4,5)P_2_) (8 %) (molar ratios), doped with Bodipy-TR ceramides at 1% molar ratio. We checked whether the lipid bilayer, deprived of septins, was fluid by performing FRAP. We obtained a diffusion coefficient of 1.36 ± 0.07 µm^2^.s^−1^ with a mobile fraction above 90 %, which is typical of lipid diffusion in a fluid membrane^32^ (supp figure 1.A). FRAP was also performed in the septin fluorescent channel, after the addition of 200 to 300 nM budding yeast GFP septins on SLBs or Giant Unilamellar Vesicles (GUVs) under polymerizing conditions (low salt). No fluorescence recovery was noticed after 10 minutes protein incubation (Figure 1A-B), suggesting that once septin filaments have assembled into filamentous structures onto membranes, they cannot diffuse anymore.

**Figure 1.**
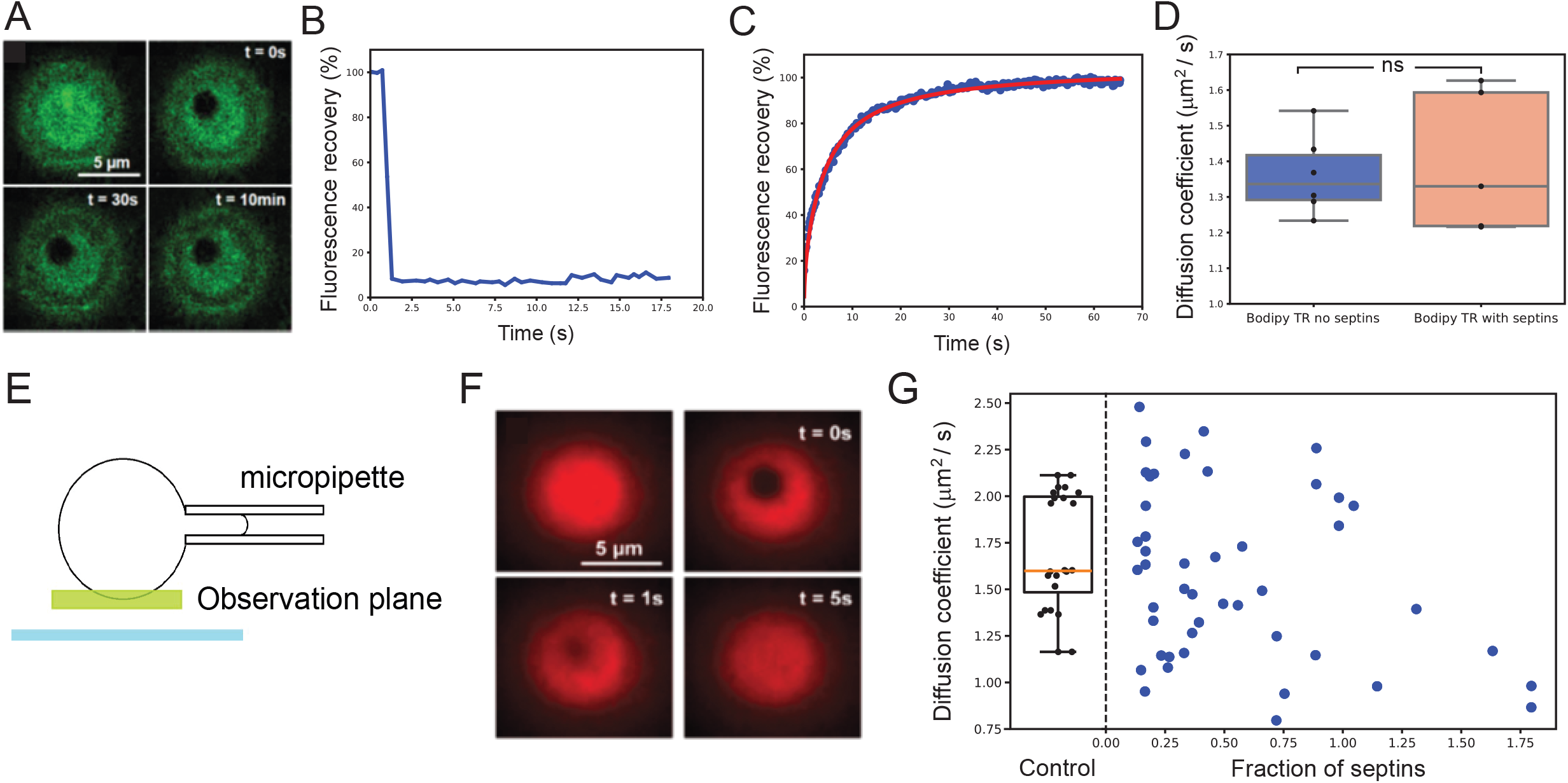
Septins minimally affect the diffusion of lipids. A. Fluorescence recovery experiment on GFP-tagged septins interacting on a GUV. Snapshots obtained with confocal microscopy at the bottom of the GUV. B. Fluorescence intensity – in percentage of the maximum intensity – in the bleached region as a function of time after correction for overall bleaching. The experiment shows no recovery in the septin signal even in the time scales of minutes. C. FRAP experiment performed on Bodipy-TR ceramide incorporated in a SLB on which septin are interacting. Fluorescence intensity in the bleached region as a function of time. D. Diffusion coefficient obtained from FRAP experiment on Bodipy-TR in control experiments without septins (left) and when septins are added (right). No significant difference was observed. E. Schematic representation of a GUV held with a micropipette, for FRAP experiment. F. Fluorescence recovery experiments performed on PI(4,5)P2 tagged with Texas Red, incorporated in GUV. Snapshots obtained using spinning disk confocal. G. Diffusion coefficient of PI(4,5)P2 in GUVs produced either by electro-formation or gel-assisted swelling. Th control experiment is shown (left) as well as the diffusion coefficient as a function of protein density (right).

We then tested whether yeast septin filaments would alter the diffusion of lipids. After incubating 300 nM of yeast septins for 30 minutes on SLBs, FRAP experiments were carried out for Bodipy-TR ceramide and resulted in a diffusion coefficient of 1.4 ± 0.12µm^2^.s^−1^ (N=3) with full fluorescence recovery, suggesting that, overall, the Bodipy-TR ceramide remained freely diffusive (Figure 1.C-D). The diffusion of lipids that do not interact with septins is thereby not affected by yeast septins.

This assay was performed, in parallel, using human septins, which self-assemble into an orthogonal network of filaments, when bound to membranes. Similarly, the diffusion of Bodipy-TR ceramide lipids from the bilayer remains largely unaffected in the presence of septins (see suppl. Figure 1.B, (N=8). We observe full fluorescence recovery following FRAP experiments, with diffusion coefficient at 2.95±0.27 µm^2^.s^−1^ in the presence of human septins and 2.9±0.18 µm^2^.s^−1^ in their absence. We notice a significant difference comparing the diffusion coefficients of lipids obtained in the presence of budding yeast versus human septins. This may result from discrepancies in the surface properties of the glass slides as experiments were performed on different days. Such variability may also explain why diffusion coefficients measured in different laboratories can differ substantially^33^.

We then wondered whether the diffusion of PI(4,5)P_2_, known to interact with septins^17^, would be affected by the presence of septins. We used commercially available Texas red PI(4,5)P_2_, in which the fluorescent probe is bound to one of the lipid tails, thereby preserving the ability of PI(4,5)P_2_ headgroup to interact with septins. The lipid mix was identical to that used above, except 6% of unlabeled PI(4,5)P_2_ was doped with 2% of TR-PI(4,5)P_2_. Strikingly, in the control SLBs, TR-PI(4,5)P_2_ was barely mobile, with a mobile fraction of 50 % only (see supp fig 1.D-E). This limited mobility is most likely due to an interaction between PI(4,5)P_2_ and the glass surface that we reported recently^34^. To bypass any artefact induced by the interaction between the lipids and a surface, we performed FRAP experiments, focusing at the bottom of a GUV held with a micropipette within a chamber (and Figure 1.E.). The spherical geometry of the membrane was accounted for in the analysis pipeline using a dedicated routine to accurately estimate and correct for distances on the curved surface, (See Suppl. Figure 2). We thus obtain a control diffusion coefficient of D = 1.7 ± 0.1 µm^2^.s^−1^ (N=3 experiments, n= 14 Guvs) for labeled PI(4,5)P_2_ (See Figure 1.G). Following the incubation with yeast GFP-septins, (N=17 experiments, n=66 Guvs) the diffusion coefficient was found to be, D = 1.6 ± 0.1 µm^2^.s^−1^ on average, suggesting that the diffusion of PI(4,5)P_2_ is barely affected by septins. In addition, the septin density – estimated by the surface fraction covered by septins (from 0 to 1.8) was systematically varied (see Figure 1.G). The septin density was estimated from a prior calibration of septin fluorescence intensity. The surface fraction, measured from the fluorescence intensity, goes beyond 1 most likely because septins have a tendency to form bundles and several layers from the membrane surface^18,35^. As shown in Figure 1.G, a slight decrease in the diffusion coefficient is observed with increasing septin density. However, this drop from about 1.6 to 1.2 µm^2^.s^−1^ is not significant. Importantly, the mobile PI(4,5)P_2_ fraction remains close to 100 % for all tested septin densities, suggesting that all lipids are still able to diffuse albeit with a slight mobility reduction. We also tested, in parallel, the role of human septins (200 nM) on the diffusion of PI(4,5). We observed, a slight decrease of diffusion coefficient (supp. Fig. 1.D) from 1.71±0.3 to 1.46±0.40 µm^2^.s^−1^ in the presence of human septins, which remains non-significant. Control experiments (without septins) with diffusion coefficients of 1.6 and 1.7 um^2^.s^−1^ for yeast and human septins, respectively were conducted using different batches of GUVs.

Taken together, our observations show that septin filaments bound to reconstituted membranes minimally impact the diffusion of lipids.

Next, we investigated whether the mobility of membrane-embedded or membrane-bound objects could be altered by septins. We used model objects containing either pure transmembrane domains or cytosolic domains of varying dimensions, up to about 30 nm in diameter. To follow their diffusion, we performed single particle tracking (SPT) experiments. Briefly, the trajectories of fluorescently labeled objects, sparsely distributed on the membrane, were tracked by optical microscopy at the bottom surface of a GUV. GUVs were held with a micropipette above a coverslip to prevent any contact between the lipid membrane and the surface. Analyzing the membrane fluorescence signal, we checked that the fluorescent vesicles were unilamellar. Since the diffusion occurred on a curved membrane, the two-dimensional displacements were corrected for the spherical geometry of the GUV (suppl Fig 2). A schematic representation of the setup is presented in Figure 1.E and an example of a single particle trajectory is shown in Suppl. Fig. 2.A.

### Free diffusion of a model transmembrane domain in a septin network

First, we labelled aquaporin, a 6 nm-diameter tetrameric water channel that lacks cytosolic domains. Aquaporin can be fully embedded within GUVs, and was labeled with Atto647N for fluorescence tracking (Figure 2.A). To avoid spectral overlap and fluorescence leakage into the aquaporin channel, the membrane was counterstained with DPPE-Atto532. We then performed single particle tracking of labelled aquaporins to follow their diffusion over a total duration of 20 seconds, with a frame exposure time of 10 ms (n=20 GUVs). Figure 2.B shows a representative snapshot of the aquaporin signal at a given time point, overlaid with the projected trajectories of individual particles. A cumulative image integrating all the time points from a control experiment is shown in Figure 2.C, highlighting the areas explored by aquaporin during the acquisition. The uniform spatial distribution of recorded positions confirms that the fluorescent probes homogeneously sample the entire membrane surface. In the absence of septins, the diffusion coefficient of aquaporin was found to be 2.4±0.0014 µm^2^.s^−1^ (N = 6 experiments, n= 24 GUVs). Upon septin incubation (200 nM), the average diffusion coefficient was only reduced to 2±0.0018 µm^2^.s^−1^ (N= 6 experiments, n= 16 GUVs) (Figure 2.D). Nonetheless, the distribution of diffusion coefficient measurements, shown in Figure 2.D, shows an increase of the immobile fraction of aquaporin in the presence of septins –15 ± 3% compared to 2 ± 0.4 % without septins. Indeed, the higher number of trajectories with diffusion coefficients laying between 10^−3^ and 10^−1^ µm^2^.s^−1^ indicate a rather restricted mobility in the presence of septins. It is possible that the immobile fraction of aquaporin originates from the presence of defects on the membrane surface (septin bundles, membrane deformations, invaginations or tubulations) that may arrest the diffusion of aquaporins.

**Figure 2.**
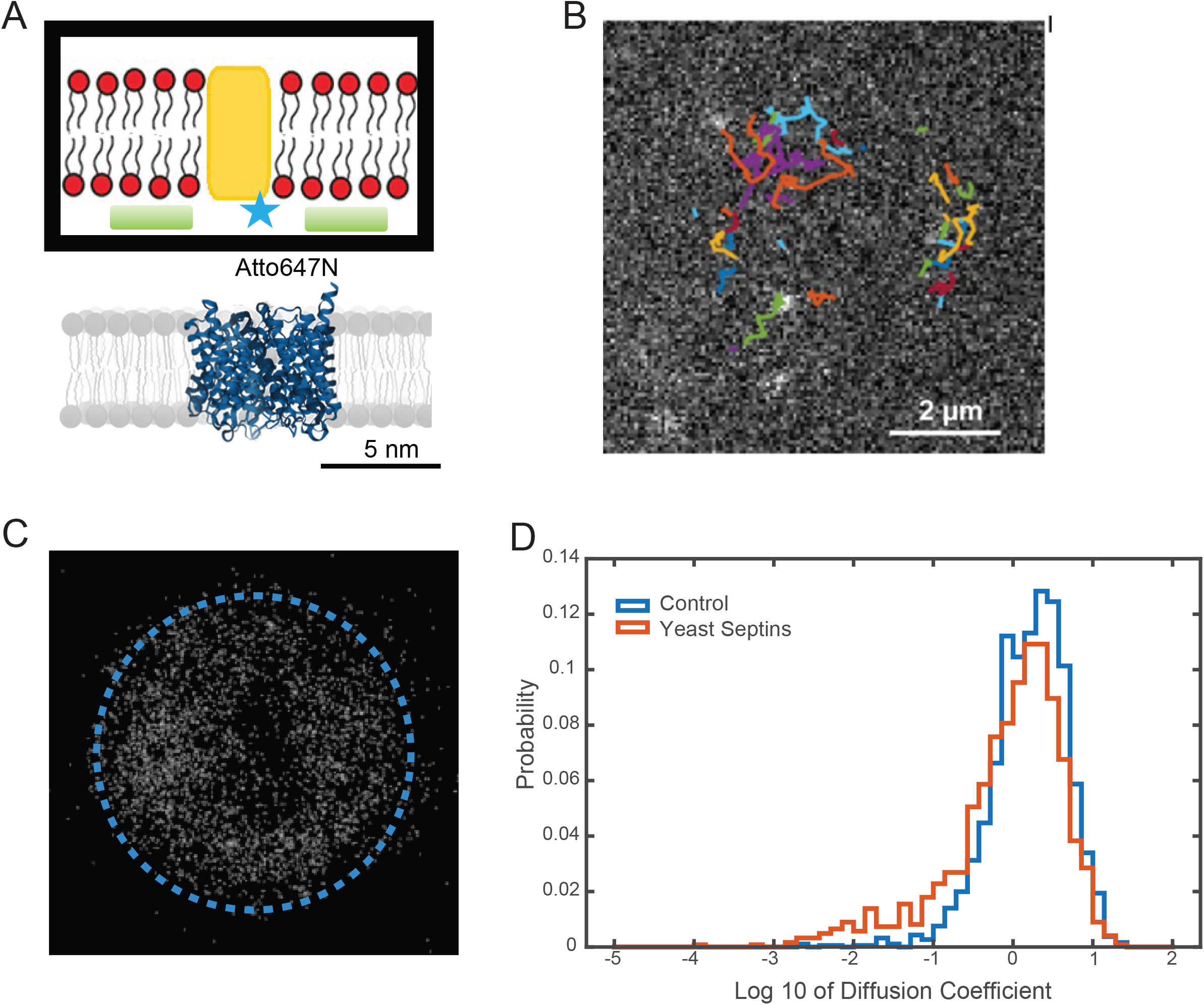
Free diffusion of a model transmembrane domain in a septin network. A. Schematic representation of aquaporin within a lipid bilayer. B. Snapshot of aquaporins at the bottom of a GUV. The exposure time is 10 ms. Trajectories obtained from the visible aquaporins are overlapped with the snapshot. Different trajectories are colour coded to distinguish them. C. Accumulation of localisations of Single Particles during a 20 s movie. On each frame all the detections are recorded, and the image is created after accumulation of all the detections (single dots) throughout the movie. D. Distribution of the diffusion coefficient of single trajectories in the control experiment and in the presence of septins. The mean diffusion coefficient is not affected but for a population of trajectories with very low diffusion coefficient – lower than 0.1 μm^2^/s.

However, because GUVs are held with a micropipette and no membrane tension is applied, we do not observe any macroscopic or marked deformations induced by septins on GUVs^19,21^. Taken together, those observations suggest that septins have a minor effect on the diffusion of aquaporins transmembrane protein.

### The diffusion of large objects is constrained by septins

In situ, septins constrain the mobility of large protein complexes bound to the membrane (polarisome^8^, exocyst complex^2^). To test whether the size of cytosolic domains influence this effect, we designed and tested the diffusion of model membrane-bound objects with cytosolic protrusions ranging from 7 to 17 nm in radius. First, to model large objects bound to the membrane we used quantum dots (QD655) passivated with streptavidin and anchored to the outer leaflet of the membrane using biotinylated lipids (0.004% of DSPE-PEG-biotin in the lipid mix). This strategy generates a membrane bound component of about 15 nm in radius, adding up the dot and the streptavidin coating. Due to the higher brightness of quantum dots compared to Atto molecules, we reduced the exposure time to 5 ms and performed SPT experiments for a total duration of 20 seconds. Septins were incubated at 200 nM with GUVs pre-decorated with Qdots after GUVs electroformation. The diffusion coefficient obtained for the control experiment (without budding yeast septins) was 2±1.4 µm^2^.s^−1^ (N = 3 experiments, n= 9 GUVs) whereas in the presence of septins, it dropped nearly tenfold to D = 0.22±0.5 µm^2^.s^−1^ (N = 3 experiments, n= 11 GUVs). Importantly, freely diffusing Qdots in solution with similar size, are known to diffuse with a diffusion coefficient around 20 µm^2^.s^−1^, thereby 100 folds larger than when bound to membranes^36^. Hence, the hindrance induced by the Qdots themselves can be considered negligible. The distributions of diffusion coefficients for both the control and septins conditions are shown in Figure 3.A. With septins bound, this distribution displays a pronounced tail towards slow-diffusing lipid-bound QDots. To estimate the fraction of slow diffusing Qdots, we set a threshold at 0.1 µm^2^.s^−1^ to define nearly immobile Qdots. Only 1.3 ± 0.2 % of Qdots were immobile in the control experiment while this proportion increased to 55 ± 7 % in the presence of septins (Figure 3.B). These results demonstrate that septins significantly hinder the mobility of Qdots bound to membrane lipids.

**Figure 3.**
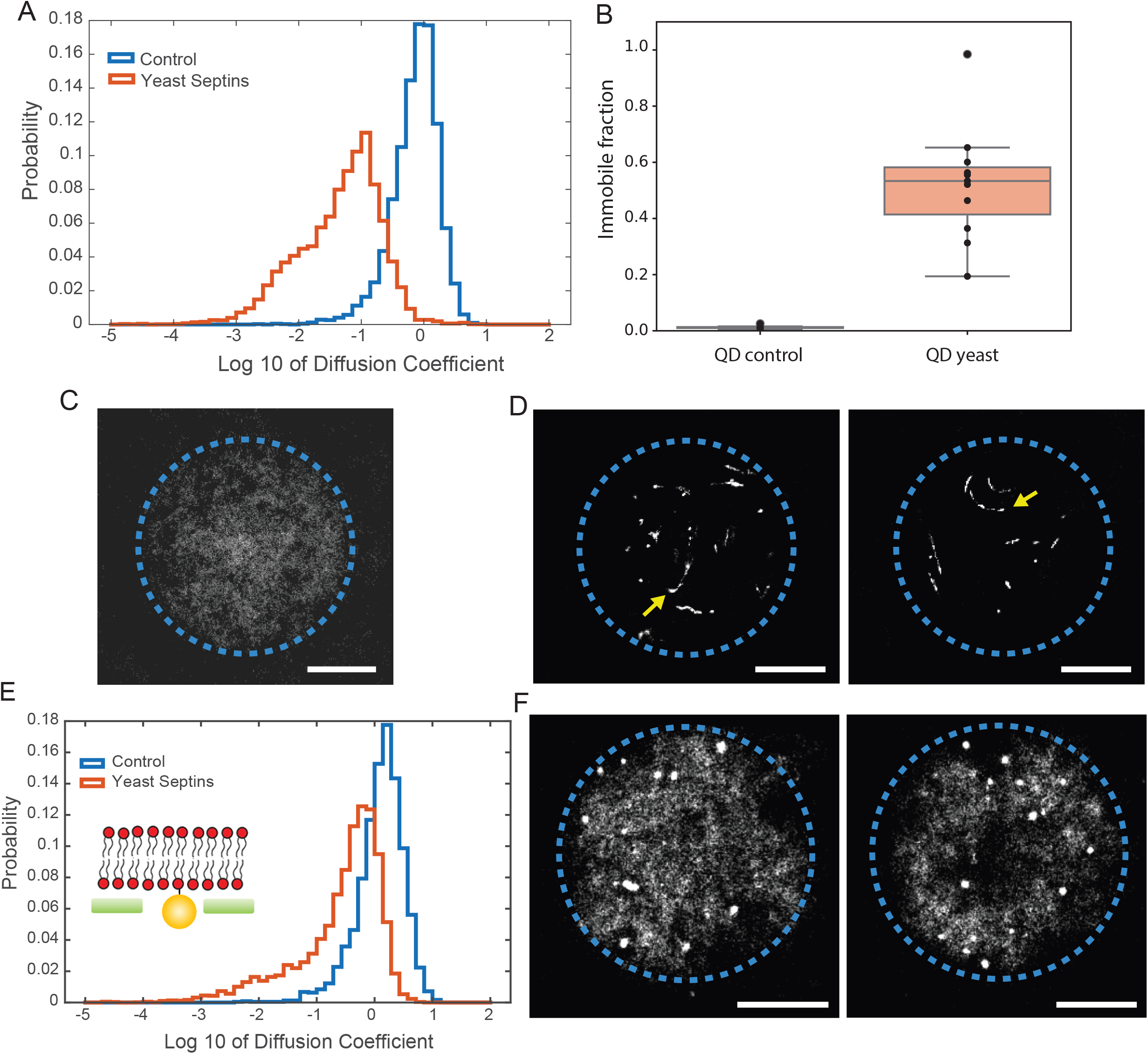
The diffusion of large objects is constrained by septins. A. Distribution of diffusion coefficients in log scale of quantum dots in the control experiment and when yeast septins interact on the membrane. B. Immobile fraction of quantum dots in the control experiment and when septins interact on the membrane. Accumulations of quantum dots during 20s movies from the control experiment (C) and after the addition of septins (D). Long linear traces (arrows) correspond to a single quantum dots diffusing in a linear region during the movie. E. Distribution of diffusion coefficients obtained for streptavidin in a logarithmic scale in the control experiment and with yeast septins bound on the vesicle. F. Accumulation of detections during 20s movies. Scale bars: 2 µm.

To further visualize how lipid-bound Qdots diffuse over the course of an experiment, we display in Figure 3.C and D the accumulation of Qdots position over time. In the control experiment, Qdots positions are homogeneously distributed in space, whereas in the presence of septins, they appear spatially confined along linear tracks of approximately 1 to 2 µm in length (pointed by arrows in figure 3.D). Budding yeast septins are known to form parallel sheets of filaments when bound to membranes^19^. Defects or gaps in this organization may create sufficient room for Qdots to move between bundled septin filaments.

Taken together, our results show that membrane-bound objects lacking cytosolic domain, such as AQP0, can freely diffuse in the presence of septin. In contrast, objects bearing large extracytosolic domains, such as30 nm lipid-anchored QDots, become trapped within the septin mesh.

We next asked whether the mobility of smaller extracytosolic domains tethered to the membrane would also be affected by septin filaments. We have therefore examined the diffusion of streptavidin molecules bound to biotinylated lipids (0.001 % in the lipid mixture). We tested that streptavidin does not bind to GUVs in the absence of biotin and in the presence of septins and was thereby not interacting with septins. Streptavidin assembles as a tetramer and extends approximately 12 nm from the membrane. Streptavidin was fluorescently labeled with Atto647N, and a bulk concentration of 20pM was found to be optimal for single particle tracking experiments. The membrane was fluorescently labeled using DPPE Atto532.

In the control experiment (in the absence of septins), the diffusion coefficient was D=1.8±0.0006 µm^2^.s^−1^ (N = 2 experiments, n=6 GUVs, see blue curve Figure 3.E), similar to that obtained for large Qdots. In the presence of septins, the distribution of diffusion coefficients (Figure 3.E, red curve) revealed a long tail towards immobile particles, with an average diffusion coefficient of D=0.56±0.00008 µm^2^.s^−1^ (N = 3 experiments, n = 9 GUVs). Even after removing the slowest trajectories with D < 0.1 µm^2^.s^−1^, the averaged diffusion coefficient remains low at D=0.71 µm^2^.s^−1^, indicating that the mobility of streptavidin molecules is significantly constrained in the presence of septins.

To further characterize the diffusion, we display the accumulation of positions of streptavidin molecules bound to lipids over the 20s acquisition duration(Figure 3.F). The control experiment, essentially similar to the observations from Figure 3.C, shows that membrane-bound streptavidin can diffuse freely on the surface of GUVs. In the presence of septins, two populations emerged: One diffusing freely and another significant remaining essentially immobile (see dense points in Figure 3.F). The quantitative analysis showed that the immobile fraction (threshold set at 0.1 µm^2^.s^−1^) increased from 2 ± 0.2 % in the control experiment to 40 ± 7% in the presence of septins.

Hence, a significant proportion of streptavidin becomes fully immobilized in the presence of septins. This immobilization could result from local defects in the lipid membrane or within the septin network. In addition, we cannot exclude the possibility that streptavidin tetramers undergo further oligomerization, forming larger assemblies that become trapped in a septin filamentous array. To assess whether the GFP tag fused to Cdc10 septin subunits influences streptavidin diffusion, we repeated the assay using dark (unlabeled) septins. In this condition, the immobile fraction of streptavidin decreased from 40% to 27±6%, while the overall distribution of diffusion coefficients remained largely similar (See supp. Figure 3). Even though this difference is not significant, it suggests that GFP tags may slightly alter septin organization leading to increased immobilization of streptavidins.

As an alternative model object, we have also analyzed the diffusion of a DSPE lipid bound to a PEG 2k and labeled with a Cy5 fluorescent probe (DSPE-PEG2k-Cy5). This object mimics a membrane-anchored protein carrying a cytosolic domain of about 8.5 nm in diameter, comparable in size to the aforementioned streptavidin probe. The radius of gyration of PEG2k is about 3.7 nm and the −Cy5 dye adds less than 1 nm in length.

We performed FRAP experiments (see supp figure 4.A) at the bottom of GUVs and observed that the diffusion of DSPE-PEG2k-Cy5 decreased in the presence of septins, with a diffusion coefficient of 1.55 ± 0.15 µm^2^.s^−1^ (N=3 experiments, n= 12 GUVs) compared to 2.4 ± 0.2 µm^2^.s^−1^ (N=2 experiments, n=7 GUVs) in the control experiment without septins. Even though the diffusion coefficient decreased with increasing septin density (See supp Figure 4.B), the mobile fraction remains at 100% across the different conditions. This thus suggests that septins slow the diffusion of membrane-bound objects with a 8.5 nm cytosolic domain, but fail to fully immobilize them.

We next asked whether the observed mobility restriction could be correlated with differences in membrane tensions. To induce high tension, GUVs were subjected to a 20% hypo-osmotic shock. For low tensions, GUVs were not held with a micropipette allowing thus the membrane to freely fluctuate and deform. The diffusion coefficients of streptavidin was found to be 0.3±0.3 µm^2^.s^−1^ (N = 4 experiments, n=12 GUVs) and 0.18±0.27 µm^2^.s^−1^ (N = 3 experiments, n=6 GUVs) for low and high tensions, respectively. The corresponding immobile fractions were 36 ±7 % for low tension and 53 ± 7 % for high tension (see supp Figure 5). These observations suggest that high membrane tension slightly reduces the mobility of streptavidins. A possible explanation is that higher tensions reduce the “membrane ruffling” and membrane fluctuations that would otherwise facilitate the passage of streptavidins across septin filaments.

**Figure 4.**
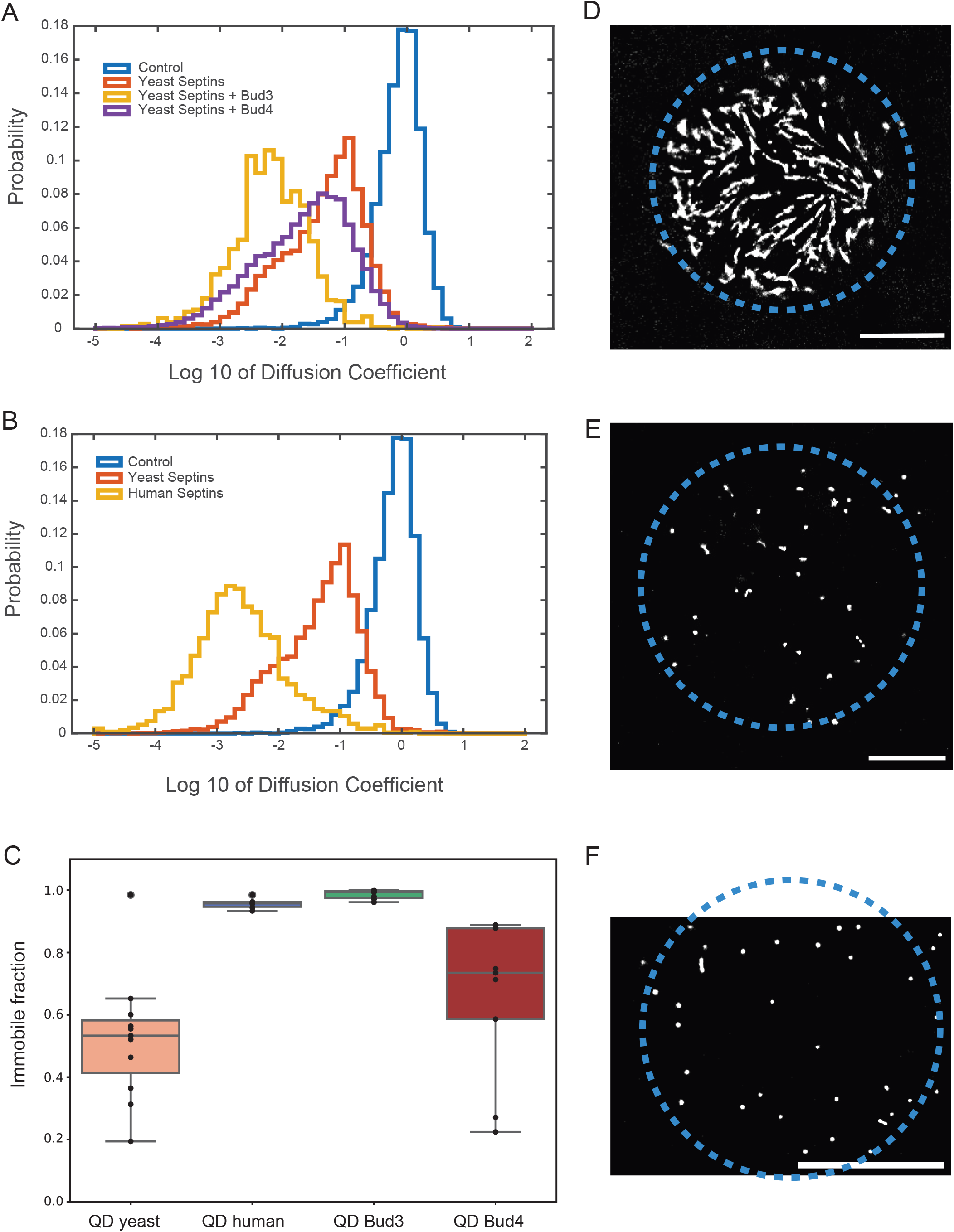
Role of the septins ultrastructural organization on diffusion. (A) and (B) Distribution of diffusion coefficient of single quantum dots in log scale for the control experiment, yeast septins bound to the membrane and different crossed-linked septin networks. (C) Immobile fraction of quantum dots for different crossed linked septin network compared to the case of yeast septins alone. (D, E and F) Accumulation of detections of quantum dots along 20s movies for respectively Bud4 + yeast setpins, Bud3 + yeast septins, and human septins. (D) Red circles correspond to +1/2 nematic defect, red arrow points at a −1/2 nematic defect.

**Figure 5.**
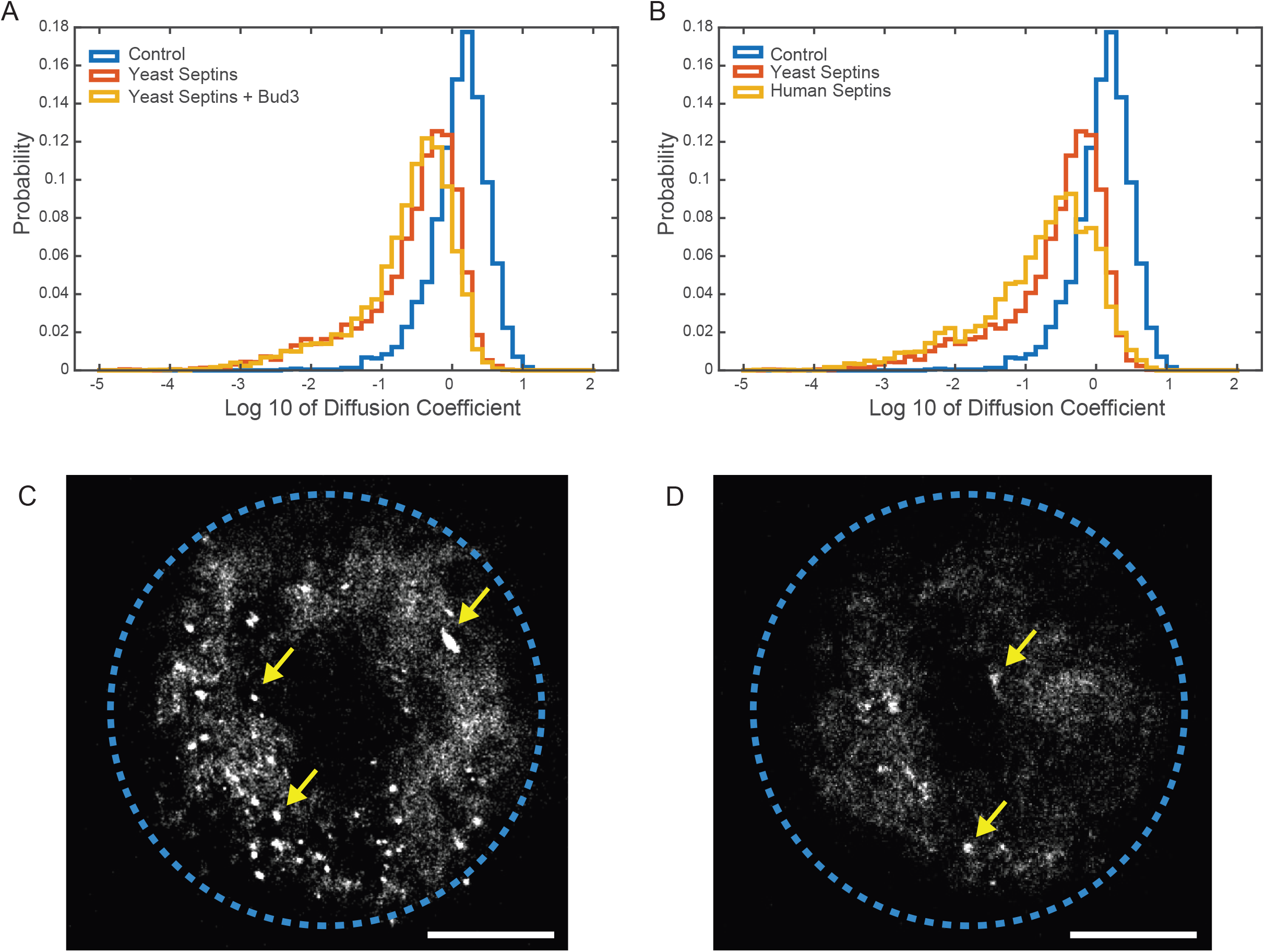
Diffusion of streptavidin in different septin networks. Single Particle Tracking of streptavidin in the presence of different cross-linked septin networks. (A and B) Distribution of diffusion coefficient of single streptavidins in log scale for the control experiment, yeast septins bound to the membrane and different crossed-linked septin networks. (C and D) Accumulation of detections of quantum dots along 20s movies for respectively Bud3 + yeast septins and human septins.

Taken together, these experiments demonstrate that the ability of septins to induce membrane diffusion barriers strongly depends on the dimensions of the diffusing object and can be modulated by membrane tension.

### Role of the septins ultrastructural organization on diffusion

The organization of septins on membranes differs depending on their nature. Budding yeast and Drosophila septin complexes were shown to assemble into parallel arrays or sheets of filaments on membranes^19,20^. In contrast, human septins tend to systematically self-assemble into orthogonal arrays of filaments on both supported lipid bilayers and liposomes^21^. Given these differences in ultrastructural organization, we wondered whether the nature of septins could impact the diffusion of the model membrane-bound objects used in this study. We therefore analyzed the role of human septins on the diffusion and mobility of these objects on membranes.

As examples, we show orthogonal arrays of human septins incubated with reconstituted membranes, liposomes, lipid monolayers or supported lipid bilayers (suppl. Figure 6.A-C). At concentrations above 40-50 nM in the bulk, human septins consistently form two-layered orthogonal filamentous networks. Interestingly, in budding yeast, specific factors appear necessary to achieve similar orthogonal organization^26^. Indeed, knocking down Bud3 and Bud4 induce a full or partial loss, respectively, of the orthogonal septin array at the hourglass^37^. To analyze any direct role of Bud3 and Bud4 in budding yeast septins organization, we performed electron microscopy experiments on purified budding yeast septins in the presence of either proteins (see suppl Figure 6.D-F). Both Bud3 and Bud4 (Supl. Fig.6.E) promoted septin bundling, while Bud3, in particular, frequently induced the formation of orthogonal arrays of filaments, resembling those observed with human septins (Suppl. Fig. 6.G and ^38^).

We next performed single particle tracking experiments within orthogonal septins arrays, induced by either human septins or by budding yeast septins in association with Bud3. In the presence of Bud3 or Bud4, we reduced septin concentrations to 100 nM, as the presence of Bud proteins enhances septin-membrane interaction and promotes filament bundling, which, in turn, interfered with SPT analysis. We choose to follow the diffusion of membrane-bound Qdots as they exhibit the most pronounced diffusion restriction in the presence of septin filaments.

The distributions of diffusion coefficients for Qdots are shown in Figure 4 for different proteins combination. When budding yeast septins were combined with Bud3 (40 nM), the average diffusion coefficient was measured at D = 0.035 µm^2^.s^−1^ (N = 3 experiments, n = 9 GUVs) indicating rather immobile Qdots. This is illustrated by of the accumulated image of all the Qdots positions over the 20s total acquisition time of the experiment, showing immobilization over time (Figure 4). With Bud4 (40 nM) and budding yeast septins, the average diffusion coefficient of Qdots D = 0.14 ± 0.25 µm^2^.s^−1^ (N = 2 experiments, n = 7 GUVs) was found to be slightly lower than with budding yeast septins alone (D = 0.22 µm^2^.s^−1^). The accumulated trajectory maps revealed that Qdots remained mobile but were constrained to linear paths, as observed for septins alone (Figure 4.D).

We compared those results with the diffusion of Qdots within an orthogonal array formed by human septins. We obtain an average diffusion coefficient of D=0.031 µm^2^.s^−1^ (N = 3 experiments, n = 7 GUVs) (see distribution in Figure 4). The accumulated snapshots images during SPT experiments, show that Qdots remain still over the course of the experiment.

The immobile fraction of Qdots was estimated to be 100% for both human septins and yeast septins in the presence of Bud3, confirming that Qdots were effectively trapped. In contrast, for yeast septins combined with Bud4, the immobile fraction reached 71±10%, which is significantly higher than for yeast septins alone (55 ± 7%). This suggests that Bud4 enhances, though to a lesser extent than Bud3, the ability of yeast septins to hinder membrane diffusion. We then asked whether the mobility of smaller model objects would also be affected by an orthogonal array of septins. We thus analyzed the diffusion of streptavidin bound to biotinylated lipids in the presence of either human septins or budding yeast septins in combination with Bud3. We obtain similar diffusion coefficients: D = 0.47 ± 0.15 µm^2^.s^−1^ (N = 4 experiments, n = 17 GUVs) for yeast septins with bud3 and 0.46± 0.14 µm^2^.s^−1^ (N = 5 experiments, n = 12 GUVs) for human septins Figure 5. The accumulation of snapshots during SPT shows a similar pattern to this observed with budding yeast septins only (Figure 5). Immobile streptavidin molecules are recognized as bright spots appearing on top of a homogeneous background of freely diffusing ones. The immobile streptavidin fractions were 44 ± 10 % for human septins and 28 ± 5 % for budding yeast/Bud3, respectively. After removing data from immobile particles, the diffusion coefficients were 0.60 µm^2^.s^−1^ for yeast septins/Bud3 and 0.67 µm^2^.s^−1^ for human. Those values are thereby close to the value obtained for budding yeast septins alone (0.71 µm^2^.s^−1^).

Taken together these observations demonstrate that the ultrastructural organization of septins controls the diffusion of large membrane bound model systems.

Using a systematic study, in a perfectly controlled environment with a minimal number of components, we could show how septins could tune the diffusion and mobility of membrane bound elements. We have clearly demonstrated here, for the first time, that the dimensions of diffusing objects as well as the nature of the ultrastructural organization of septins were key to tune membrane proteins diffusion.

## Discussion

In this paper, we demonstrate that an array of septin filaments assembled at the membrane can directly hinder the diffusion of membrane-associated components, providing thus evidence for a long-standing hypothesis that septins are involved in the formation of diffusion barriers. We used in vitro reconstituted systems, to isolate the physical effects of septins from other cellular factors and show that their barrier function depends on the size of the extramembrane domain of the diffusing objects.

Our results reveal a clear size influence of the diffusing molecule given septins restrict the diffusion of the largest biomimetic objects we have tested. Objects of 30 nm in diameter are fully immobilized by septin, while membrane bound components of about 10 nm in diameter retain some mobility. This is consistent with in vivo observations, in which the mobility of Spa2 from the polarisome and Sec5p and Sec3p from the exocyst complex (30 nm large cytosolic domain) was controlled by a septin dependent diffusion barrier^8^.

Several factors likely contribute to this size-dependent filtering. Membrane fluctuations and thus nanometric deformations would induce ruffles that may allow small enough components to slip between septin filaments and the membrane. Indeed, we observed that osmotic shock on the membrane slightly affected the diffusive behavior of streptavidins in a septin network. Additionally, electron tomography experiments had shown that septin filaments in budding yeast can be spaced from the membrane^39^, suggesting that septin membrane anchoring may be indirect at specific time point during cell division. Such anchoring might be mediated by additional adaptor proteins, or by septin coiled coils that bridge the membrane at a distance.

Unexpectedly, we found that septins had only a minor effect on the diffusion of PI(4,5)P_2_, despite their known affinity for this lipid. This result contrasts with recent in vivo studies^15^, which reported significant restriction of the diffusion of PI(4,5)P_2_ by septins and spectrins as opposed to other proteins : ER-PM contact sites, actin or clathrin coated structures. These discrepancies may stem from multiple factors. First, the live cell experiments performed in Pacheco et al^15^. may introduce additional players, such as actin, that act synergistically with septins and arrest the diffusion of PI(4,5)P_2_ ^22,40^. Second, our results highlight that the size of the fluorescent probe plays a crucial role in determining whether septins act as effective diffusion barriers in our single-particle tracking (SPT) experiments. “Tubbyc” probes were used to optimize the PALM experiments in Pacheco and their dimension of about 10 nm may have influenced their diffusion behavior within septin assemblies. In our reconstituted system, we observe that the mobility of probes with comparable dimensions is significantly reduced in the presence of septin filaments,.

In situ, septins may act as a size-selective filter, restricting the mobility of large membrane-bound complexes while allowing smaller specific components to diffuse. In addition, the ultrastructural organization of septins, either as parallel sheet of nematically organized filaments or as orthogonal arrays at the membrane seems to be crucial to direct the nature of mobility restrictions. Septins are known to undergo dramatic structural rearrangements, and this intrinsic plasticity may serve as a regulatory mechanism to control and finely adjust the diffusion of membrane proteins or lipids at specific time points during cellular processes or at specific cellular locations.

For example, we have shown here a key role for Bud3 and Bud4, two associated proteins which are necessary to ensure a proper organization of septins splitted rings towards the end of cytokinesis, in the diffusion restriction. Prior to cell spitting, these proteins contribute in maintaining essential cellular factors in between the septin splitted rings, ensuring successful membrane scission. Moreover, septins have been shown to be curvature-sensitive in various contexts^19,21,31,41,42^. This curvature sensitivity likely contributes to the spatial targeting of septin assemblies to physiologically relevant membrane sites where diffusion barriers are required. This would be for instance particularly relevant at the base of cilia, base of dendrites, annulus of spermatozoa or at the intercellular bridge during mammalian cell division. Our data suggests that budding yeast septins need additional factors (Bud3) to act as efficient diffusion barriers. This assumption was confirmed in budding yeast where Bud proteins were shown to be key for sequestrating specific proteins -Aim4, Nba1, Nis1 and Spa2-at the bud neck through budding^38^. In contract, human septins in our assay readily self-assemble into orthogonal filaments and fully prevent the diffusion of Qdot. This difference may reflect adaptation to cellular division timing between species: the time scales required for cytokinesis in yeast is much shorter than the cytokinetic process in human cells where intercellular bridges can persist for hours before final membrane fission. It is thus possible that additional proteins facilitate the plasticity and reorganization of budding yeast septins to efficiently and rapidly induce cell division.

In addition to septins, other factors can restrict the diffusion of membrane proteins or lipids. One contributor is the high concentration of membrane proteins in situ, which can account for up to 50% of the total mass in cell membranes – much higher than the levels used here to optimize SPT assays. Under such conditions, molecular crowding alone can significantly reduce the mobility of membrane-bound objects. Furthermore, membrane contact sites, such as those between PM and ER can locally hinder the diffusion of proteins. Lipid composition also plays a role; certain lipids, such as long-chain sphingolipids and ceramides^10^, which accumulate at specific membrane domains, have been shown to be key in inducing steric diffusion barriers. Taken together, our findings alongside other reports suggest that septins alone are not selective diffusion barriers. Instead, they likely act in concert with other cellular component to efficiently restrict the mobility of membrane-bound proteins.

Taken together, our observations establish a direct and steric role for septins in the formation of diffusion barriers at the membrane. In vitro reconstituted assays, as carried out here, efficiently highlight the specificity of septins in this process. In vitro bottom-up assays thereby finely dissect the respective role and specific function of each components in complex cellular functions. Future work using multi-component reconstituted systems with enhanced complexity intermes of substrate geometries so we can test various curvatures and additional protein partners will be essential to unravel how septins collaborate with other elements to fine-tune membrane diffusion barriers.

## Methods

### Reagents and material

Common reagents (ethanol, acetone, chloroform, sucrose, sodium chloride, Tris) were purchased from VWR reagents and Sigma-Aldrich Co.. L-α-phosphatidylcholine (EggPC, 840051P), cholesterol (700000P), 1,2-dioleoyl-sn-glycero-3-phosphoethanolamine (DOPE, 850725P), 1,2-dioleoyl-sn-glycero-3-phospho-L-serine (DOPS, 840035P), and L-α-phosphatidylinositol-4,5-bisphosphate (PI(4,5)P2, 840046P) were purchased from Avanti polar lipids. Bodipy-TR-ceramide was purchased from Invitrogen (D-7540). Atto532, Atto647N, Atto647Nand DPPE Atto532 were purchased from Sigma-Aldrich. QD655 was purchased from Thermo Fischer scientific.

### Protein expression and purification

Bud3 and Bud4 proteins were gifts from the Piatti lab (CRBM, Montpellier).

Budding yeast septins were purified according to the established protocols in Bertin et al.^43^. Cdc10-GFP-Cdc3-Cdc12-Cdc11 complexes were expressed in E.Coli before being subjected to Nickel affinity, Size exclusion and ion exchange purification. Human septins were purified as describes in Iv et al^44^.. Briefly, septins are purified by nickel-histidine, streptavidin-biotin affinities and size exclusion chromatography. Septin complexes consist of octamers SEPT2GFP-SEPT6-SEPT7-SEPT9-SEPT9-SEPT7-SEPT6-SEPT2GFP. Those complexes are stored in high ionic strength solutions (300mM KCl, 50mM Tris pH 8) at −80 °C (after quick freeze in liquid nitrogen) to avoid polymerization and aggregation of complexes in solution.

### Preparation of SLBs

Supported lipid bilayers were generated by the fusion of SUVs (Small Unilamellar Vesicles) on a hydrophilic substrate. Various protocols have been optimized for the formation of SLBs, and were summarized by Lin et al^45^. SUVs were prepared by mixing 0.5 mga lipids mix (EggPC 56.5%, Cholesterol 15%, DOPE 10%, DOPS 10%, PI(4,5)P2 8% and Bodipy TR ceramide 0.5% or other fluorescent lipids). The mix dissolved in Choroform was dried under gentle nitrogen flow before being placed under vacuum for 30 min to remove any trace of solvent. The lipid film was then rehydrated in 125 μL of “SUV Buffer” (150 mM NaCl and 20 mM Citrate (pH = 4.8)) for 30 min. The solution was vortexed for a few seconds until it became opaque and then sonicated in a water bath sonicator (Elmasonic s10) for 5 to 10 minutes until it became transparent again. The SUVs were then aliquoted and could be stored for up to 4 weeks at −20°C. A 25 µL aliquot of SUVs was thawed for each experiment and diluted in 125 μL of “SUV Buffer” to reach a final lipid concentration of 0.7 mg/mL. The obtained solution was added in the chambers and left to incubate at room temperature for 30 min. The chambers were then washed thoroughly by pipetting in in out 5 times using the “SUV Buffer” and additional 5 times using the “Observation Buffer”. The washing steps are crucial for obtaining good bilayers. The obtained bilayers were used immediately. When septins were added, they were left to incubate for 30 min.

### Preparation of GUVs

The “Growth Buffer” (50 mM NaCl, 50 mM Sucrose and 10 mM Tris (pH = 7.8)) and the “Observation Buffer” (75 mM NaCl and 10 mM Tris (pH=7.8)) were prepared. Their osmolarities were controlled with a freezing point osmometer (Löser) and adjusted by adding small amounts of NaCl until the osmolarity difference was below 5%. Giant Unilamellar Vesicles (GUVs) were prepared by either electro-formation from platinum wires or PVA-gel assisted methods with lipid mixtures of EggPC 56.5%, Cholesterol 15%, DOPE 10%, DOPS 10%, PI(4,5)P2 8% and Bodipy TR ceramide 0.5% or other fluorescent lipids^17^. In both methods, lipids were resuspended in 10 mM Tris pH 7.8, 50mM NaCl and sucrose. GUVs were collected and transferred in solution with 10mM Tris pH 7.8 and a given NaCl concentration. Osmolarity inside and outside of vesicles were adjusted with sucrose according to the NaCl concentration of the external solution.

### Preparation of Proteo-Guvs holding aquaporins

Aquaporin 0 (AQP0) was purified from bovine eye lenses. The bovine lenses were flash frozen in liquid nitrogen and stored at −80°C. After slow thawing at 4°C in 10 mM Tris (pH = 8), 5 mM EDTA, 5 mM EGTA and a protease inhibitor mix, the nucleus and the cortex were separated. The cortex was then grinded with Potter-Elvehjem into small pieces. The obtained solution was then washed, ultra centrifuged (45 min at 55,000 rpm) and resuspended in a succession of different buffers, first : 10 mM Tris (pH = 8) 4 mM Urea, second : 10 mM Tris (pH = 8) 20 mM NaOH, third : 10 mM Tris (pH = 8). The solution was solubilized in 4% (w/w) Octyl β-D-Glucopyranoside (OG) for 3h at 4 °C. After ultra centrifugation (20 min at 200,000 g) the supernatant was added to an ion exchange mono-S column (Amersham Biosciences) pre equilibrated with Tris 10 mM (pH = 8) 1.5% OG. The protein was eluted with a gradient from 0% to 100 % of buffer containing Tris 10 mM (pH = 8) 350 mM NaCl 1.5 % OG. Fractions containing aquaporin were pooled together and injected onto a size exclusion column Superose 12 /60.

To incorporate the aquaporins into SUVs, detergent was added at a concentration between “Rsat”, corresponding to the beginning of the SUVs solubilization and “Rsol” corresponding to the SUVs full solubilization and the micellar state. Octyl β-DGlucopyranoside at a concentration of 40 mM was used. Aquaporins were then added with a weight ratio of proteins to lipids of 1:100. The solution was left at 4°C for 1 hour. To remove detergent, a total weight ratio of 20/1 Biobeads (Bio-Rad)/detergent was used^46^. They were first cleaned using methanol and rinsed with MilliQ water. The addition of Biobeads was performed in three steps with 1h incubation at 4°C. After each step, the supernatant was collected and the beads were discarded. Finally, the proteo-SUVs solution was aliquoted, flash frozen and kept at −80°C for further use.

### Fluorescence Recovery after Photobleaching (FRAP)

FRAP experiments were performed using both a spinning disk confocal microscope and a TIRF microscope. The FRAP module was controlled using the ILAS2 software combined with Metamorph. Different durations were used to sample the recovery of the fluorescence. Directly after the laser burst, the time intervals were set to detect the fast kinetic of the initial recovery. It was later increased by a factor 5 to obtain the final kinetics of the recovery. The maximum laser intensity was used for bleaching to shorten the time needed to fully bleach the region of interest (ROI). The laser was focused in the observation plane and scanned within the region of interest to uniformly photo-bleach the molecules. The number of scans required was optimized to achieve full bleaching with minimal time. Typically, bleaching times of 200-300 ms were used. Bleaching times of 50 to 200 ms were used respectively for GUVs or SLBs. The analysis was performed using a custom made Matlab routine to correct for the background fluorescence as well as the bleaching of the whole sample, due to the exposure for observation. Bleaching caused by the observation can be corrected using the fluorescence intensity of a region far enough and not affected by the laser burst.

The theory for analyzing FRAP recovery curves has been thoroughly. The most simple solution and still often used, is to fit the data with an exponential decay function. In this study, the bleached area is assumed to have been homogeneously bleached, and therefore have homogeneous initial fluorescence intensity. In this case the fluorescence as a function of time goes as :

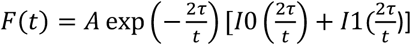

Where I0 and I1 are the modified Bessel functions, A is the mobile fraction and τ is the characteristic time of the recovery. It is directly linked to the diffusion coefficient by :

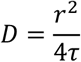

Where D is the diffusion coefficient, and r is the radius of the disk where bleaching occurs.

### Single particle tracking

A Nikon TIEclipse microscope equipped with a TIRF arm from Nikon was used. It possessed three laser lines (λ = 488 nm; λ = 532 nm and λ = 638 nm). The lasers were controlled with a Digital to Analog Converter (DAC). The calibration of the laser power output as a function of the applied voltage was performed. The Prime95B (Photometrics) camera was used for single particle detection. 4 different labeled particles were tracked: DSPE-PEG-Cy5, aquaporins labelled with Atto647, streptavidin-Atto647N and Quantum Dots. We used a X100 objective with an aperture number of 1.49. Data was acquired with Metamorph.

The glass slides used for the chambers were cleaned by sonication with acetone, ethanol and water each step for 10 min before being dried under a flow of pressurized nitrogen with a filter to prevent oil deposition on the glass coverslips. A later plasma cleaning procedure was sometimes used.

The density of particles was optimized by testing a range of particle concentrations in the membrane. The concentration used for Quantum Dots was 50 pM and 20 pM for streptavidin. The molar ratio of DSPE-PEG2k-biotin was optimized at 0.004% for Quantum Dots and 0.002% for streptavidin. Both were incubated with vesicles after electro-formation for 1 hour at 4°C under gentle mixing. Using this protocol only the lipids of the external leaflet of the GUVs were labelled. Therefore the fluorescent particles were facing directly towards potential septins, interacting. The vesicles where then transferred in the observation buffer where they would be incubated with septins. GUVs were held using a micropipette. Images were collected every 5 to 10 ms and the final acquisition time was 20 s.

### Single particle tracking: data analysis and correction for sphericity

Acquired movies were analyzed using a custom made Matlab script “SlimFast” to obtain the localization and to assign them to trajectories. To compute the diffusion coefficient of the trajectories, the mean squared displacement (MSD) was computed as a function of the time step. Considering Brownian motion and particles diffusing in 2 dimensions, the apparent diffusion coefficient was extracted based on MSD(t) = 4Dt. Where D is the diffusion coefficient, t the time step used to compute MSD. To fit experimental data, one has to consider effects coming from the imperfect precision of localization as well as the motion blur^47^. The formula then becomes :

MSD(t) = 4Dt + 4(σ2 − 2DΔtR), Where the second term accounts for the precision of localization σ and the motion blur R. A low number of points are actually needed for optimal fitting to compute the apparent diffusion coefficient assuming pure Brownian and to avoid noises at higher time lags. The first five points of the curve (from 1dt to 5 dt) were chosen to give a good fit to the model.

## Acknowledgements

We thank Patricia Bassereau for useful advice and discussions. We thank Manos Mavrakis for the kind gifts of the plasmids expressing human septins. This work benefited from the support of the ANR (Agence Nationale de la Recherche) for funding the project “SEPTIME”, ANR-13-JSV8-0002-01 and the project “SEPTSCORT”, ANR-20-CE11-0014-01. B. Chauvin is funded by the Ecole Doctorale “ED564: Physique en Ile de France” and Fondation pour la Recherche Médicale. K. Nakazawa was supported by Sorbonne Université (AAP Emergence). We thank the Labex Cell(n)Scale (ANR-11-LABX0038) and to Paris Sciences et Lettres (ANR-10-IDEX-0001-02). We thank the Cell and Tissue Imaging (PICT-IBiSA), Institut Curie, member of the French National Research Infrastructure France-BioImaging (ANR10-INBS-04).

## Author contributions

A.B, B.H., S.M. and B.C. designed research. A.B., K.N. and B.C. expressed and purified the septin complexes. S.P and I.A. expressed and purified Bud proteins. B.C., K.N. and B.H. performed the fluorescence microscopy experiments. B.C. and B.H. analyzed the diffusion data. The manuscript was written by A.B, S.M., B.H. and B.C. The results interpretation were discussed by all of the authors.

## References

1. Caudron, F. & Barral, Y. Septins and the lateral compartmentalization of eukaryotic membranes. Dev Cell 16, 493–506 (2009).

2. Dobbelaere, J. & Barral, Y. Spatial Coordination of Cytokinetic Events by Compartmentalization of the Cell Cortex. Science 305, 393–396 (2004).

3. Longtine, M. S. et al. Septin-dependent assembly of a cell cycle-regulatory module in Saccharomyces cerevisiae. Mol Cell Biol 20, 4049–4061 (2000).

4. DeMarini, D. J. et al. A septin-based hierarchy of proteins required for localized deposition of chitin in the Saccharomyces cerevisiae cell wall. J Cell Biol 139, 75–93 (1997).

5. Hu, Q. et al. A septin diffusion barrier at the base of the primary cilium maintains ciliary membrane protein distribution. Science 329, 436–439 (2010).

6. Ihara, M. et al. Cortical organization by the septin cytoskeleton is essential for structural and mechanical integrity of mammalian spermatozoa. Dev Cell 8, 343–352 (2005).

7. Ewers, H. et al. A Septin-Dependent Diffusion Barrier at Dendritic Spine Necks. PLoS One 9, e113916 (2014).

8. Barral, Y., Mermall, V., Mooseker, M. S. & Snyder, M. Compartmentalization of the cell cortex by septins is required for maintenance of cell polarity in yeast. Mol Cell 5, 841–851 (2000).

9. Kwitny, S., Klaus, A. V. & Hunnicutt, G. R. The annulus of the mouse sperm tail is required to establish a membrane diffusion barrier that is engaged during the late steps of spermiogenesis. Biol Reprod 82, 669–678 (2010).

10. Clay, L. et al. A sphingolipid-dependent diffusion barrier confines ER stress to the yeast mother cell. Elife 3, e01883 (2014).

11. Fujiwara, T. Phospholipids undergo hop diffusion in compartmentalized cell membrane. The Journal of Cell Biology 157, 1071–1082 (2002).

12. Gowrishankar, K. et al. Active remodeling of cortical actin regulates spatiotemporal organization of cell surface molecules. Cell 149, 1353–1367 (2012).

13. Jaqaman, K. et al. Cytoskeletal control of CD36 diffusion promotes its receptor and signaling function. Cell 146, 593–606 (2011).

14. Heinemann, F., Vogel, S. K. & Schwille, P. Lateral membrane diffusion modulated by a minimal actin cortex. Biophys J 104, 1465–1475 (2013).

15. Pacheco, J., Cassidy, A. C., Zewe, J. P., Wills, R. C. & Hammond, G. R. V. PI(4,5)P2 diffuses freely in the plasma membrane even within high-density effector protein complexes. J Cell Biol 222, (2023).

16. Zhang, J. et al. Phosphatidylinositol polyphosphate binding to the mammalian septin H5 is modulated by GTP. Curr Biol 9, 1458–1467 (1999).

17. Beber, A. et al. Septin-based readout of PI(4,5)P2 incorporation into membranes of giant unilamellar vesicles. Cytoskeleton (Hoboken) 76, 92–103 (2019).

18. Bertin, A. et al. Phosphatidylinositol-4,5-bisphosphate promotes budding yeast septin filament assembly and organization. J Mol Biol 404, 711–731 (2010).

19. Beber, A. et al. Membrane reshaping by micrometric curvature sensitive septin filaments. Nat Commun 10, 420 (2019).

20. Szuba, A. et al. Membrane binding controls ordered self-assembly of animal septins. Elife 10, (2021).

21. Nakazawa, K. et al. A human septin octamer complex sensitive to membrane curvature drives membrane deformation with a specific mesh-like organization. J Cell Sci 136, (2023).

22. Mavrakis, M. et al. Septins promote F-actin ring formation by crosslinking actin filaments into curved bundles. Nat Cell Biol 16, 322–334 (2014).

23. Kuzmić, M. et al. Septin-microtubule association via a motif unique to isoform 1 of septin 9 tunes stress fibers. J Cell Sci 135, jcs258850 (2022).

24. Spiliotis, E. T. Regulation of microtubule organization and functions by septin GTPases. Cytoskeleton (Hoboken) 67, 339–345 (2010).

25. Sadian, Y. et al. The role of Cdc42 and Gic1 in the regulation of septin filament formation and dissociation. Elife 2, e01085 (2013).

26. Chen, X., Wang, K., Svitkina, T. & Bi, E. Critical Roles of a RhoGEF-Anillin Module in Septin Architectural Remodeling during Cytokinesis. Curr Biol 30, 1477–1490.e3 (2020).

27. Wloka, C. et al. Evidence that a septin diffusion barrier is dispensable for cytokinesis in budding yeast. Biol Chem 392, 813–829 (2011).

28. Wu, H., Guo, J., Zhou, Y.-T. & Gao, X.-D. The anillin-related region of Bud4 is the major functional determinant for Bud4’s function in septin organization during bud growth and axial bud site selection in budding yeast. Eukaryot Cell 14, 241–251 (2015).

29. Kang, P. J., Hood-DeGrenier, J. K. & Park, H.-O. Coupling of septins to the axial landmark by Bud4 in budding yeast. J Cell Sci 126, 1218–1226 (2013).

30. Bridges, A. A. et al. Septin assemblies form by diffusion-driven annealing on membranes. Proc Natl Acad Sci U S A 111, 2146–2151 (2014).

31. Cannon, K. S., Woods, B. L., Crutchley, J. M. & Gladfelter, A. S. An amphipathic helix enables septins to sense micrometer-scale membrane curvature. J Cell Biol 218, 1128–1137 (2019).

32. Lindblom, G. & Orädd, G. Lipid lateral diffusion and membrane heterogeneity. Biochim Biophys Acta 1788, 234–244 (2009).

33. Rose, M., Hirmiz, N., Moran-Mirabal, J. M. & Fradin, C. Lipid Diffusion in Supported Lipid Bilayers: A Comparison between Line-Scanning Fluorescence Correlation Spectroscopy and Single-Particle Tracking. Membranes (Basel) 5, 702–721 (2015).

34. Vial, A. et al. Correlative AFM and fluorescence imaging demonstrate nanoscale membrane remodeling and ring-like and tubular structure formation by septins. Nanoscale 13, 12484–12493 (2021).

35. Taveneau, C., Blanc, R., Péhau-Arnaudet, G., Di Cicco, A. & Bertin, A. Synergistic role of nucleotides and lipids for the self-assembly of Shs1 septin oligomers. Biochem J 477, 2697–2714 (2020).

36. McHale, K., Berglund, A. J. & Mabuchi, H. Quantum dot photon statistics measured by three-dimensional particle tracking. Nano Lett 7, 3535–3539 (2007).

37. Ong, K., Svitkina, T. & Bi, E. Visualization of in vivo septin ultrastructures by platinum replica electron microscopy. Methods Cell Biol 136, 73–97 (2016).

38. Adriaans, I. E. et al. Architecture and function of Bud3 and Bud4-induced septin structures. bioRxiv 2025.10.03.680245 (2025) doi:10.1101/2025.10.03.680245.

39. Bertin, A. et al. Three-dimensional ultrastructure of the septin filament network in Saccharomyces cerevisiae. Mol Biol Cell 23, 423–432 (2012).

40. Kinoshita, M., Field, C. M., Coughlin, M. L., Straight, A. F. & Mitchison, T. J. Self- and actin-templated assembly of Mammalian septins. Dev Cell 3, 791–802 (2002).

41. Bridges, A. A., Jentzsch, M. S., Oakes, P. W., Occhipinti, P. & Gladfelter, A. S. Micron-scale plasma membrane curvature is recognized by the septin cytoskeleton. J Cell Biol 213, 23–32 (2016).

42. Lobato-Márquez, D. et al. Mechanistic insight into bacterial entrapment by septin cage reconstitution. Nat Commun 12, 4511 (2021).

43. Bertin, A. et al. Saccharomyces cerevisiae septins: Supramolecular organization of heterooligomers and the mechanism of filament assembly. PNAS 105, 8274–8279 (2008).

44. Iv, F. et al. Insights into animal septins using recombinant human septin octamers with distinct SEPT9 isoforms. J Cell Sci 134, jcs258484 (2021).

45. Lin, W.-C., Yu, C.-H., Triffo, S. & Groves, J. T. Supported membrane formation, characterization, functionalization, and patterning for application in biological science and technology. Curr Protoc Chem Biol 2, 235–269 (2010).

46. Rigaud, J. L., Paternostre, M. T. & Bluzat, A. Mechanisms of membrane protein insertion into liposomes during reconstitution procedures involving the use of detergents. 2. Incorporation of the light-driven proton pump bacteriorhodopsin. Biochemistry 27, 2677–88 (1988).

47. Vestergaard, C. L., Blainey, P. C. & Flyvbjerg, H. Optimal estimation of diffusion coefficients from single-particle trajectories. Phys Rev E Stat Nonlin Soft Matter Phys 89, 022726 (2014).

