## supplemental material for "Septin Architecture Dictates Size-Dependent Diffusion Barriers on biomimetic Membranes"

**Affiliations:**

### Supplementary Figure 1

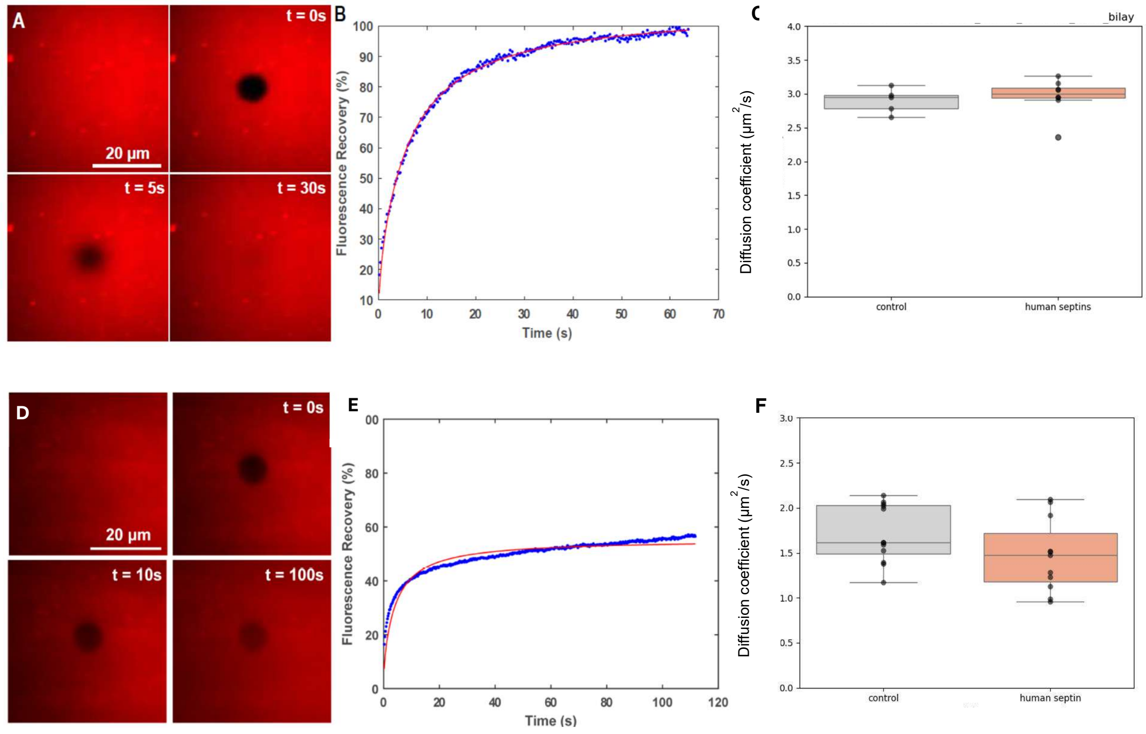


**A**. Fluorescence recovery experiment on Bodipy-TR ceramide in a lipid bilayer deposited on glass. Snapshots obtained with TIRF microscopy during the fluorescence experiment with t = 0 corresponding to the bleaching step. **B.** Fluorescence intensity – in percentage of the maximum intensity – in the bleached region as a function of time after correction for overall bleaching. It shows that the mobile fraction is high with a diffusion coefficient of DBdpy−TR = 1.36±0.07 µm2/s in good agreement with the literature. **C**. Diffusion coefficient of Bodipy-TR ceramide in a supported lipid bilayer, in the presence of human septins.

**D**. Fluorescence recovery experiments performed on PI(4,5)P2 tagged with Texas Red, incorporated in SLB deposited either on glass. Snapshots obtained with TIRF microscopy. After a few minutes, the bleached zone can still be distinguished. **E**. Fluorescence intensity of the bleached zone as a function of time after overall bleaching correction. Half of the lipids do not fully recover after photobleaching. **F**. Diffusion coefficient of Bodipy-TR ceramide in a GUV, in the presence of human septins.

### Supplementary Figure 2

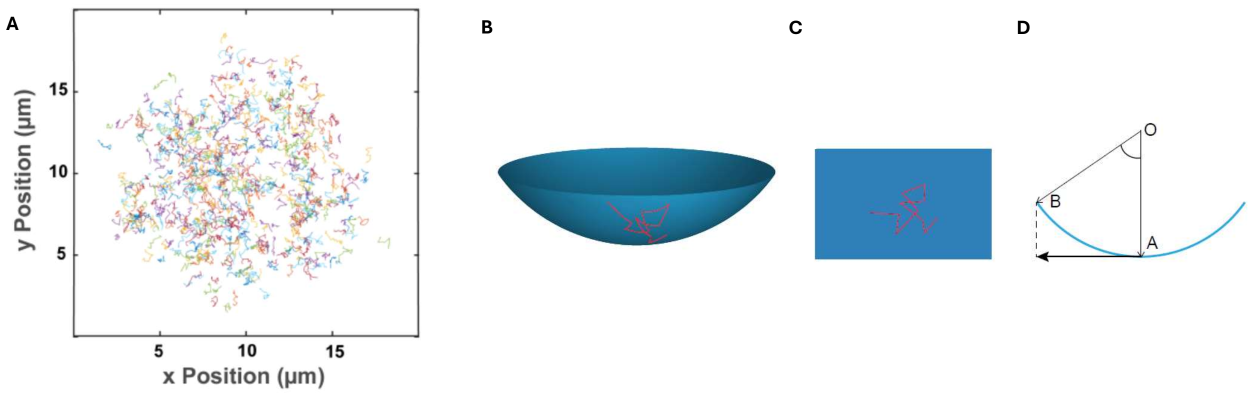


**A**. Trajectories obtained from the bottom of a GUV, with different colors to distinguish the individual trajectories. Correction for the calculation of distances for the single particle experiment. **B.** Trajectories follow the shape of the vesicle while the camera only gives the projection on a plane (**C**). The calculation for the actual distance can be performed by looking at the length of the arc between the two position (**D**).

### Supplementary Figure 3

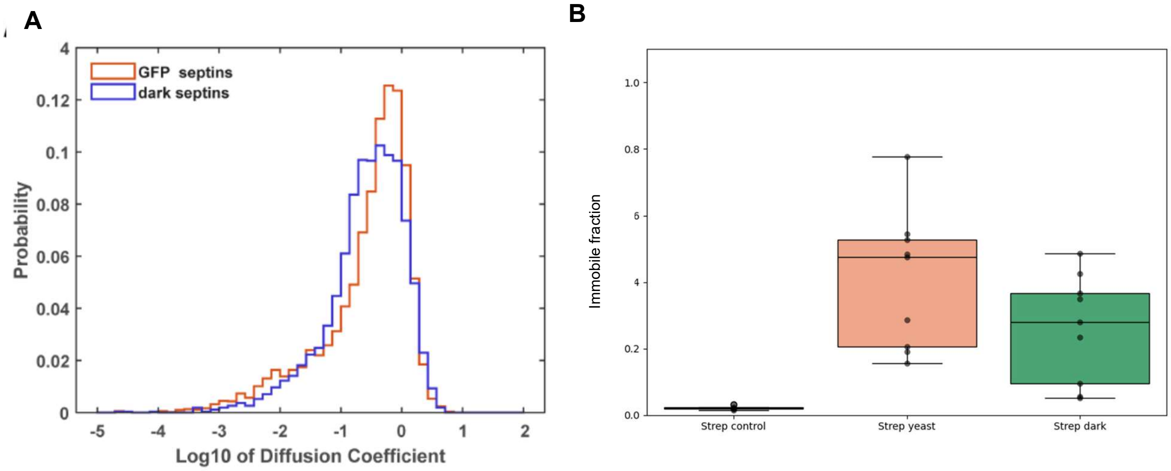


**A**. Distribution of diffusion coefficient of single trajectories of streptavidin for the GFP-tagged yeast septins and for the control with unlabelled septins. **B**. Comparison of the immobile fraction of streptavidin in the control experiment and in the presence of yeast septins, labelled or unlabelled.

**Supplementary Figure 4**

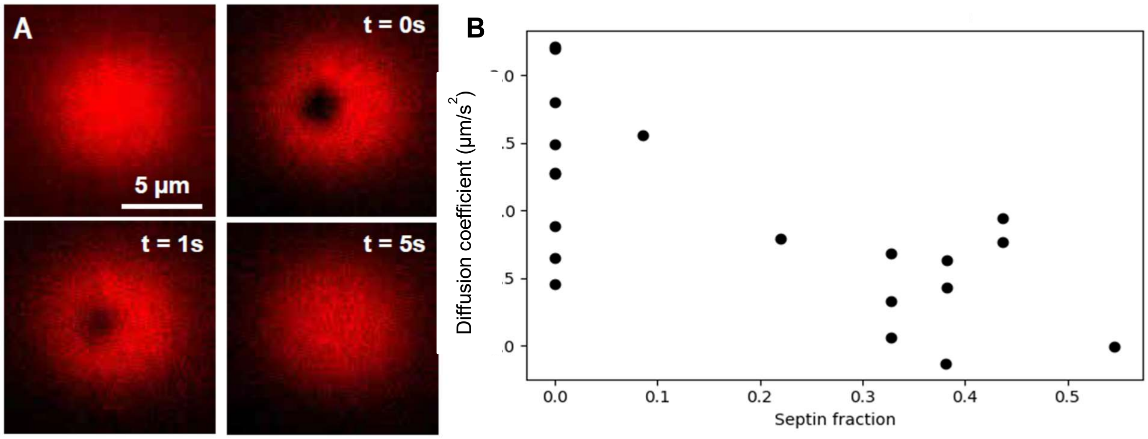


Fluorescence recovery experiment performed on DSPE-PEG2k-Cy5 lipid incorporated in a GUV produced by electro-formation. **A**. Snapshots obtained using spinning disk confocal. **B**. Diffusion coefficient in the control and as a function of the septin density. A slight drop of diffusion coefficient can be observed when septin density increases.

**Supplementary Figure 5**

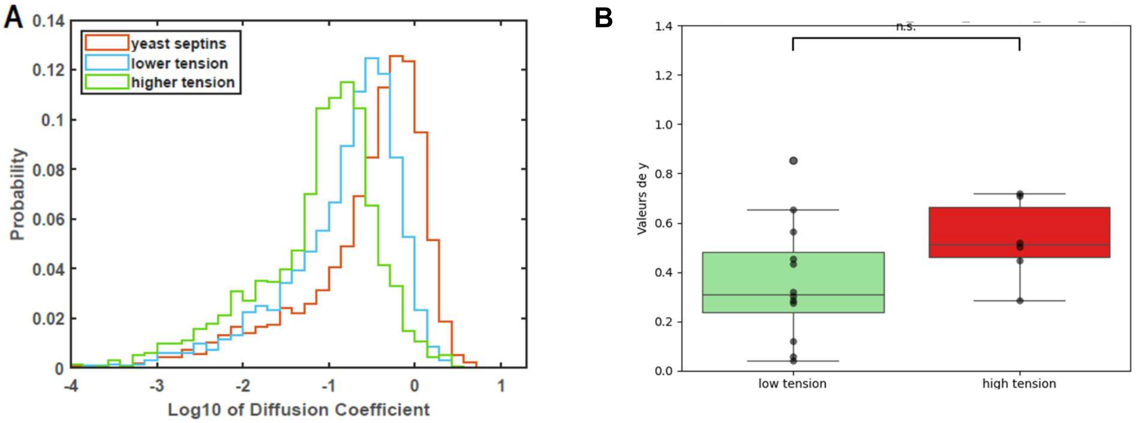


**A**. Distribution of diffusion coefficient for single trajectories of streptavidin bound to DSPE-PEG2k-biotin in the presence of yeast septins with either lower or higher tension in the membrane. **B**. Immobile fraction of streptavidin in presence of yeast septins at either lower or higher tension.

**Supplementary Figure 6**

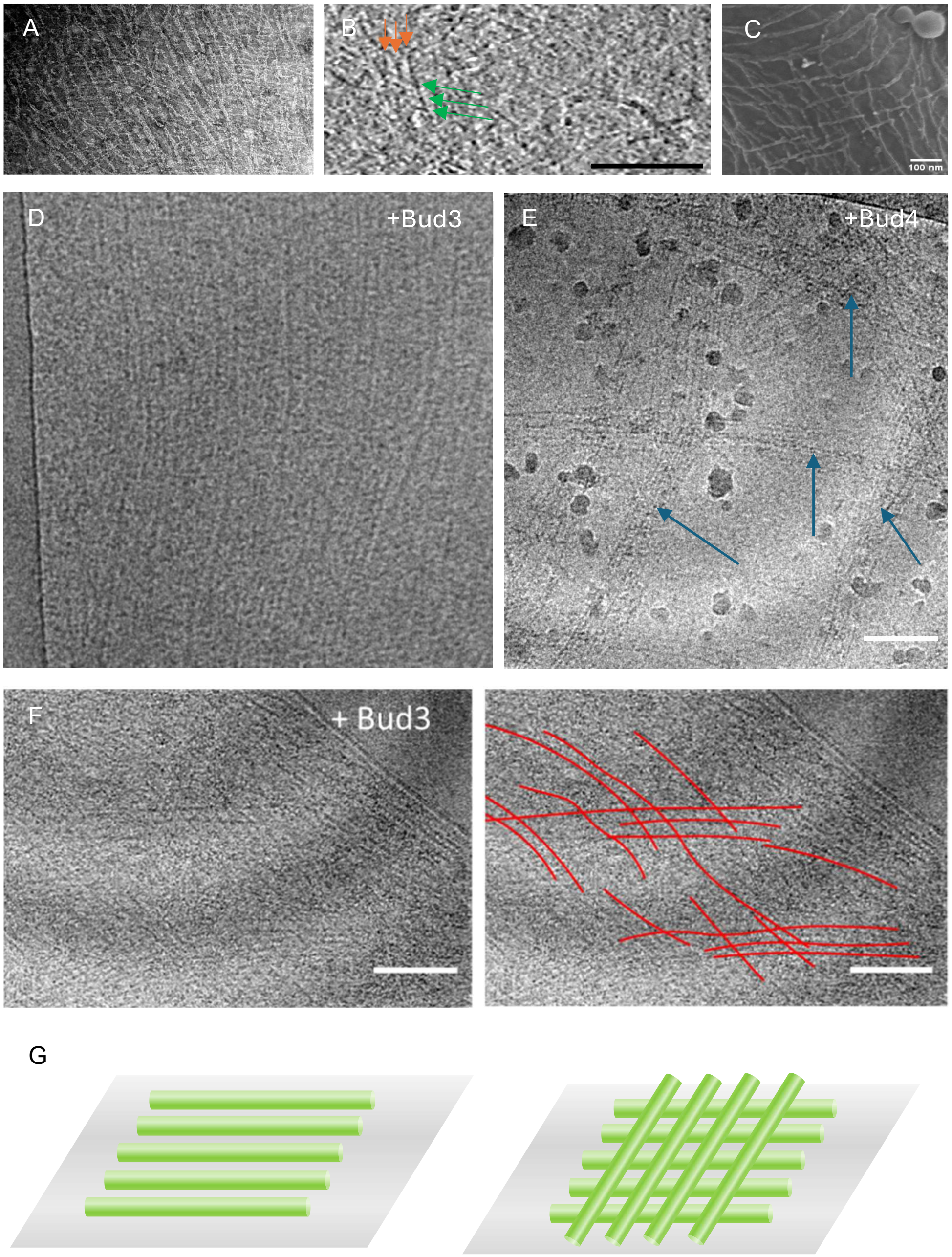


**A.** Human septins bound to a lipid monolayer and organized in an orthogonal array of filament. **B**. Slice of a cryo-tomogram displaying human septin filaments arranged orthogonally and bound to liposomes. Arrows indicate filaments' ends**. C**. Scanning Electron Microscopy image of a human septin orthogonal network bound to a SLB.

**D-F**. Images from cryo electon microscopy performed on yeast septins (100 nM) in solutions mixed respectively with Bud3 (**D**) and Bud4 (**E**) (200 nM). Bud 3 organizes yeast septins into arrays (**D-F**) while Bud4 bundles septin filaments (**E**)**. F**. Images of yeast septins and Bud3 bound to liposomes. Filaments are delineated in red. **G**. Schematic representation of septin ultrastructures as parallel (left) or orthogonal arrays (right).
